# CD1d-dependent neuroinflammation impairs tissue repair and functional recovery following a spinal cord injury

**DOI:** 10.1101/2023.10.13.562047

**Authors:** Xiangbing Wu, Jianyun Liu, Wei Li, Mohammad Faizan Khan, Heqiao Dai, Jeremy Tian, Raj Priya, Daniel J. Tian, Wei Wu, Alan Yaacoub, Jun Gu, Fahim Syed, Christopher H. Yu, Xiang Gao, Qigui Yu, Xiao-Ming Xu, Randy R. Brutkiewicz

## Abstract

Tissue damage resulting from a spinal cord injury (SCI) is primarily driven by a robust neuroimmune/neuroinflammatory response. This intricate process is mainly governed by a multitude of cytokines and cell surface proteins in the central nervous system (CNS). However, the critical components of the neuroimmune/neuroinflammatory response during SCI are still not well-defined. In this study, we investigated the impact of CD1d, an MHC class I-like molecule mostly known for presenting lipid antigens to natural killer T (NKT) cells and regulating immune/inflammatory responses, on neuroimmune/neuroinflammatory responses induced by SCI. We observed an increased expression of CD1d on various cell types within the spinal cord, including microglia/macrophages, oligodendrocytes (ODCs), and endothelial cells (DCs), but not on neurons or astrocytes post-SCI. In comparison to wildtype (WT) mice, a T10 contusive SCI in CD1d knockout (CD1dKO or *Cd1d*^-/-^) mice resulted in markedly reduced proinflammatory cytokine release, microglia/macrophage activation and proliferation. Following SCI, the levels of inflammatory cytokines and activation/proliferation of microglia/macrophages were dramatically reduced, while anti-inflammatory cytokines such as IL-4 and growth factors like VEGF were substantially increased in the spinal cord tissues of CD1dKO mice when compared to WT mice. In the post-acute phase of SCI (day 7 post-SCI), CD1dKO mice had a significantly higher frequency of tissue-repairing macrophages, but not other types of immune cells, in the injured spinal cord tissues compared to WT mice. Moreover, CD1d-deficiency protected spinal cord neuronal cells and tissue, promoting functional recovery after a SCI. However, the neuroinflammation in WT mouse spinal cords was independent of the canonical CD1d/NKT cell axis. Finally, treatment of injured mice with a CD1d-specific monoclonal antibody significantly enhanced neuroprotection and improved functional recovery. Therefore, CD1d promotes the proinflammatory response following a SCI and represents a potential therapeutic target for spinal cord repair.

**Significance Statement:** The cell surface molecule, CD1d, is known to be recognized by cells of the immune system. To our knowledge, this is the first observation that the CD1d molecule significantly contributes to neuroinflammation following a spinal cord injury (SCI) in a manner independent of the CD1d/NKT cell axis. This is important, because this work reveals CD1d as a potential therapeutic target following an acute SCI for which there are currently no effective treatments.

## Introduction

A traumatic spinal cord injury (SCI) represents a significant public health issue, affecting more than 2.5 million people worldwide with approximately 130,000 new injuries reported annually (1). SCI is a common cause of permanent disability in children and adults with a tremendous socioeconomic impact on affected individuals and the health care system (2). SCI is composed of primary and secondary injuries (2). A primary injury is caused by the initial traumatic impact, whereas the secondary injury is an indirect result of destructive immunological, inflammatory, neurotoxic, and biochemical cascades that are initially triggered by the primary injury.

Secondary injury remains an important determinant of SCI outcome as its damage can often be in excess of that caused by primary injury (2). Secondary injury begins within minutes following the initial primary injury and can be temporally divided into acute (<48 h after the traumatic event), sub-acute (48 h to 14 days), intermediate (14 days to 6 months), and chronic (> 6 months) phases (2, 3). In the clinical management of SCI, neurological outcomes are generally determined at 72 h after SCI (3, 4), indicating that acute and subacute phases represent the most important time window in which to achieve optimal tissue repair and neurological recovery. Accordingly, neuroimmune and neuroinflammatory responses play a central role in orchestrating the secondary injury mechanisms in the acute and sub-acute phases of SCI (3). During the acute and subacute phases, immune cells such as monocytes/macrophages (M/M*ɸ*), T cells, B cells, and neutrophils infiltrate the spinal cord to interact with neural cells and produce inflammatory cytokines, chemokines, and neurotoxic molecules that contribute to cell death and tissue degeneration (3). Although many immune cell types are involved in these early phases of SCI, because of their highly flexible programming (5), M/M*ɸ* have been shown to exhibit critical regulatory activity in response to environmental cues and play an active and dual role in SCI. Whereas M/M*ɸ* responses are characterized by the accumulation of inflammatory M/M*ɸ* in the acute phase to clear cellular debris, there is also the production of inflammatory cytokines/chemokines such as IL-1β, IL-6, IL-8, and TNF-α, thereby exerting both beneficial and detrimental effects (6, 7). Inflammatory M/M*ɸ* responses are tightly regulated in a timely manner. Importantly, inflammatory M/M*ɸ* can be polarized into tissue-repairing M/M*ɸ* by cytokines such as IL-4 in the local tissue microenvironment. These then produce growth factors such as vascular endothelial growth factor (VEGF) and anti-inflammatory mediators (e.g., IL-4, IL-10, and TGF-β1) to foster inflammation resolution, tissue repair, and wound healing (8, 9). In addition, M/M*ɸ* and microglia (tissue-resident macrophages that are exclusively derived from yolk sac-derived progenitors) coordinate cellular interactions with neuronal and non-neuronal cells in the spinal cord to restore tissue homeostasis after SCI (10). However, the cellular and molecular mechanisms of immune homeostasis and resolution of inflammation in the spinal cord after SCI are only partially understood. It is critical to unravel these mechanisms for developing effective treatments for this devastating condition to fill a significant unmet medical need.

A SCI leads to tissue destruction and cellular damage which, in turn, rapidly release endogenous “danger” molecules, known as damage-associated molecular patterns (DAMPs). DAMPs prime innate immune cells such as infiltrating M/M*ɸ* and neutrophils, as well as the resident microglia and astrocytes in the central nervous system (CNS) to induce sterile neuroimmune/neuroinflammatory responses (6, 7, 11). Because lipids are essential structural and functional components of all cell types (12), lipids are also released and have been identified as DAMPs (13). The CNS has a rich lipid composition with about 75% of all mammalian lipid species that are exclusively present in neural tissues (14), demonstrating the unique requirements of lipids for neural cell functions such as synaptogenesis and neurogenesis, impulse transmission, and the development and maintenance of the CNS (15, 16). The glycolipids represent a dominant class of lipids in the myelin bilayer of neural cells (17), which may act as DAMPs to directly trigger innate immune responses in the spinal cord after SCI. The released lipids may also be processed as lipid antigens that are presented by CD1d molecules to natural killer T (NKT) cells (18–20), a subset of CD1d-restricted T cells that bridges innate and adaptive immunity (21). CD1d is a major histocompatibility complex (MHC) class I-like molecule that, unlike the classical MHC class I molecules that present peptides to CD8 T cells, presents lipid antigens to NKT cells (18, 19). The CD1d/NKT (natural killer T) cell axis has been shown to be important in a variety of diseases including some CNS diseases such as multiple sclerosis (MS) and Experimental Allergic Encephalomyelitis (EAE) (22, 23). In addition to presentation of lipid antigens to NKT cells, CD1d also exerts intrinsic signaling to regulate immune and inflammatory responses (24–27). To our knowledge, neither CD1d nor NKT cells have been studied in the context of SCI.

We have analyzed the response of CD1d knockout mice (CD1dKO, which do not have NKT cells) to a contusive SCI at the 10^th^ thoracic vertebra (T10). Using a diverse array of motor and behavioral assessments, we found that CD1dKO mice recovered significantly faster and better from SCI than wildtype (WT) mice. Concomitantly, less damage to the spinal cord and more spared white matter were observed in CD1dKO mice when compared to WT mice. In addition, the levels of inflammatory cytokines and activation/proliferation of microglia/M*ɸ* were dramatically reduced, whereas anti-inflammatory cytokines (e.g., IL-4) and growth factors (e.g., VEGF) were detected at much higher amounts in spinal cord tissues of CD1dKO mice when compared to WT mice. In the subacute phase of SCI (i.e., day 7 post-SCI), CD1dKO spinal cord tissues had a significantly higher frequency of tissue-repairing M/M*ɸ* vs WT mice. Treatment of SCI mice with a CD1d-specific monoclonal antibody (mAb) significantly enhanced neuroprotection and improved functional recovery. Strikingly, these CD1d effects appear to be mediated by mechanisms independent of the canonical CD1d/NKT cell axis. Thus, CD1d promotes neuroimmune/neuroinflammatory responses following SCI and represents a potential therapeutic target for SCI.

## Results

### CD1d expression on spinal cord cells was upregulated upon a SCI

CD1d not only presents lipid antigens to NKT cells but also exerts intrinsic signaling to regulate immune and inflammatory responses (24–27). Such CD1d activities have not been studied in the CNS, including the spinal cord. To investigate whether and which spinal cord cells express CD1d, we performed immunofluorescence microscopy (IFM) analysis of primary rat spinal cord cells and immunohistochemistry (IHC) analysis of rat spinal cord tissue sections. IFM analysis showed co-expression of CD1d and Iba1 (a microglial/macrophage-specific marker), CD1d and Olig2 (oligodendrocyte transcription factor 2, a specific marker of oligodendrocyte progenitor cells or ODCs), or CD1d and CD31 (the most sensitive marker for endothelial cells or ECs; Fig. 1A, left panel), indicating that CD1d was constitutively expressed on primary rat spinal cord microglia/macrophages, ODCs, and ECs. These results are in line with previous reports (28, 29). CD1d was not present on the surface of either neurons (i.e., NeuN^+^ cells) or astrocytes (i.e., GFAP^+^ cells) (Fig. 1A, left panel). Our IHC analyses revealed the same distribution and phenotype of CD1d expression (Fig. 1A, right panel).

**Fig. 1.**
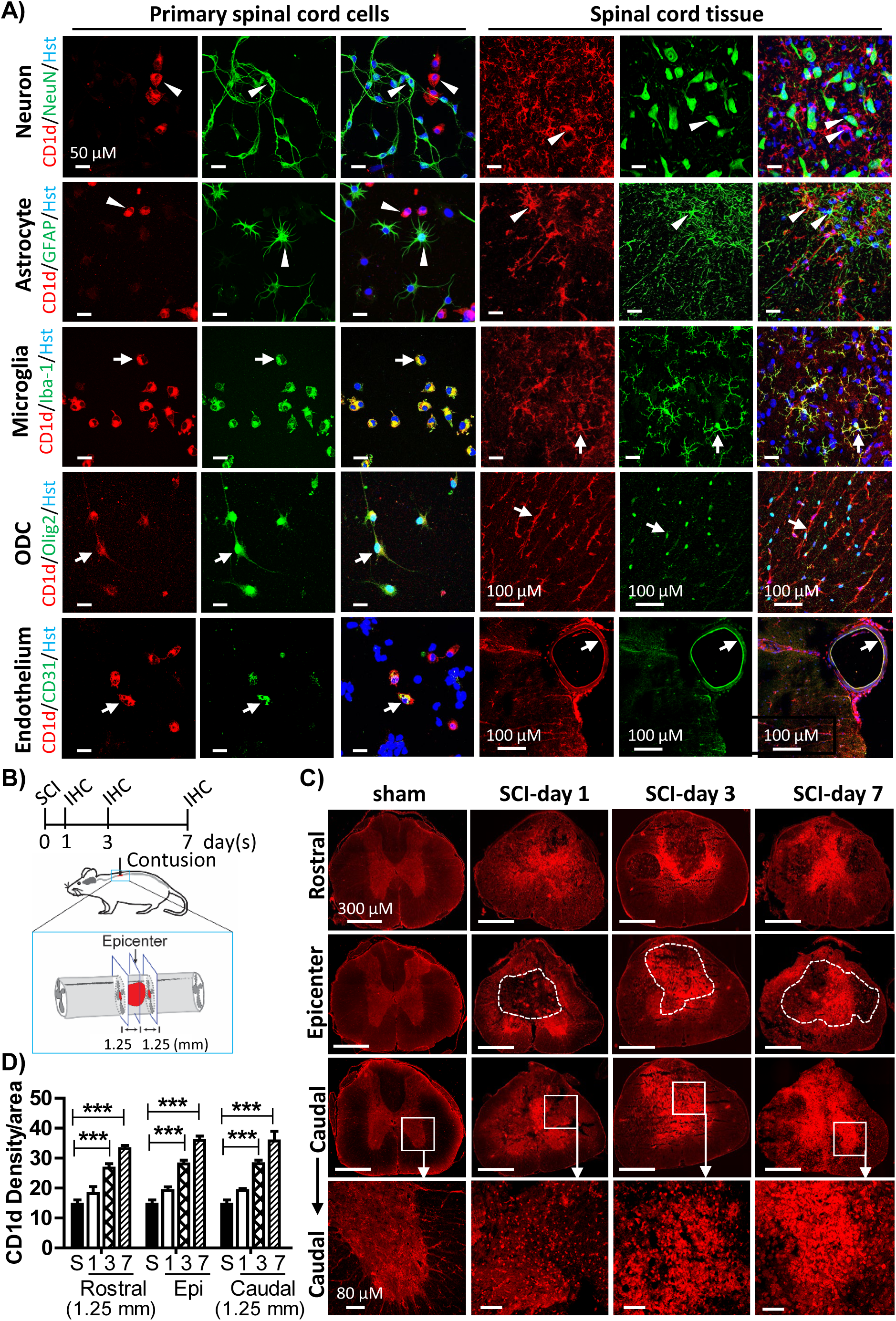
CD1d expression on spinal cord cells was upregulated upon a SCI. **A)** Left panel: IFM analysis of CD1d (red) plus either NeuN (neurons, green), GFAP (astrocytes, green), Iba1 (microglia, green), Oligo2 (ODCs, green), or CD31 (ECs, green) in primary rat spinal cord cells. Hoechst dye (Hst, blue) was for nuclear staining. Right panel: IHC analysis of CD1d expression in normal rat spinal cord tissue. Cells CD1d^+^ only or CD1d^+^Iba1^+^, CD1d^+^Olig2^+^, or CD1d^+^CD31^+^ double positive (white arrowheads); cells not co-labeled with CD1d/NeuN or CD1d/GFAP (white arrows). White bar: 50 μM. **B)** Spinal cord from days 1, 3 and 7 post-SCI vs sham-injured rats. **C)** CD1d in the injured epicenter and 1.25 mm away (rostral and caudal) 7-days post-SCI. White short and long bars: 80 μM and 300 μM, respectively. **D)** Pooled data showing CD1d expression in the injured epicenter, as well rostral and caudal to the injury site over the 7-day analysis post-SCI. Student’s t-test: \*\*\**p* < 0.001. n ≥ 3 per group. S, sham.

To investigate whether CD1d protein expression was affected by a SCI, we performed IHC analysis of CD1d in rat spinal cord tissues that were collected from rats on days 1, 3, and 7 post-SCI or rats with a sham injury (Fig. 1B). Compared to sham-injured rats, CD1d gradually and markedly increased in the injured epicenter, both rostral and caudal, over the 7-day analysis following SCI (Fig. 1C, 1D). Thus, in the spinal cord, CD1d is constitutively expressed on microglia/M*ɸ*, ODCs, and ECs, but not on neurons or astrocytes. CD1d expression is upregulated upon a SCI.

### CD1dKO mice exhibited less spinal cord damage after a SCI

Similar to the rats, increased CD1d expression over 7 days was detected in mouse spinal cord tissues following a T10 moderate contusive SCI (Fig. 2A). To analyze the effects of CD1d on histological outcomes of the spinal cords from mice post-SCI, we stained tissue sections with Cresyl Violet (also known as Nissl) and Luxol Fast Blue (LFB) to evaluate lesion area size and amount of spared white matter (SWM) (Fig. 2B), respectively. On day 7 post-SCI, in WT mice, the averages of lesion size and SWM volume were 41.1% and 35.2%, respectively (Fig. 2C). In CD1dKO mice, the averages of lesion size and SWM volume were 23.8% and 53.3%, respectively (Fig. 2C). Thus, spinal cords of CD1dKO mice had less damage and greater amounts of SWM than WT mice after injury. We also generated a 3D reconstruction of the injured spinal cord areas based on Nissl and LFB staining data (Fig. 2D). As is evident, compared to WT mice, injured spinal cords from CD1dKO mice had a smaller lesion volume and a shorter length of spinal cord damage, and a greater volume of SWM after SCI (Fig. 2D, 2E). Thus, CD1dKO mice exhibit less spinal cord tissue damage than WT mice after a SCI.

**Fig. 2.**
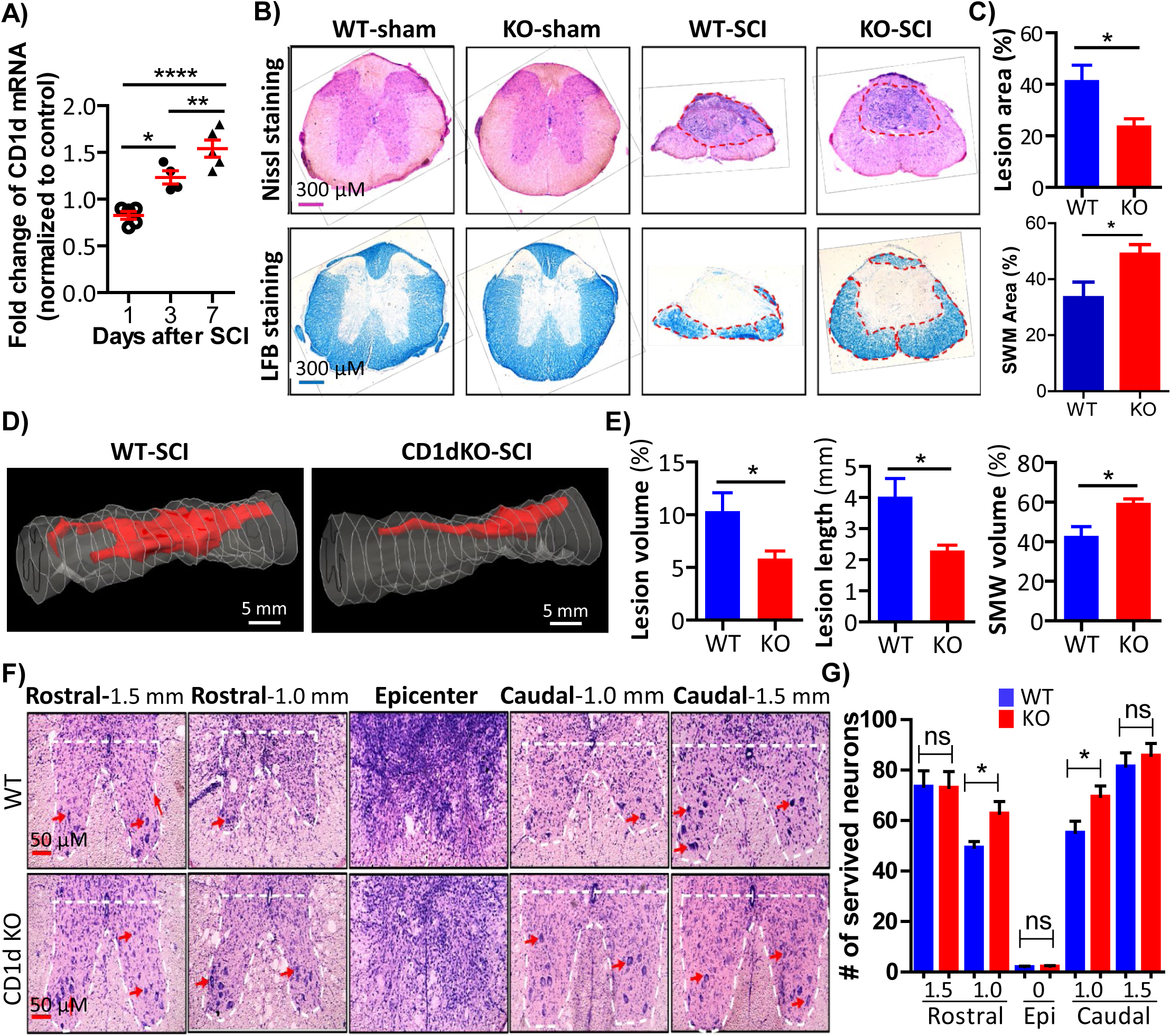
SCI-triggered tissue damage in spinal cords from CD1dKO vs WT mice. **A)** qRT-PCR analysis of WT mouse spinal cord CD1d mRNA levels over 7-days post-SCI. **B)** Representative images of spinal cord Nissl (upper panel) and Luxol Fast Blue (LFB) staining (lower panel) showing lesion area size and spared white matter (SWM), respectively. **C)** Pooled data of lesion area size and SWM from WT (blue bars) and CD1dKO (red bars) mice. **D)** 3D reconstruction of the injured spinal cord areas based on Nissl and LFB staining data from WT and CD1dKO mice. **E)** Pooled data of 3D reconstruction results (blue bars: WT mice; red bars: CD1dKO mice). **F)** Representative images of IHC analysis of viable spinal cord motor neurons (red arrows) in CD1dKO vs WT mice post-SCI using Nissl staining. G) Pooled data of viable spinal cord motor neurons in CD1dKO (red bars) vs WT (blue bars) mice post-SCI from Nissl staining. Student’s t-test: \**p*<0.05; ns, not significant. WT-SCI, n=12; CD1dKO-SCI, n=13; WT-sham, n=6; CD1dKO-sham, n=6.

With less spinal cord damage in CD1dKO mice post-SCI, we speculated that viable motor neurons might show enhanced protection. We performed IHC analysis of viable motor neurons in the spinal cords of CD1dKO vs WT mice post-SCI. There were almost no viable motor neurons in the ventral horns at the injury epicenters of both WT and CD1dKO mice post-SCI (Fig. 2F, 2G). However, in marked contrast, 1.0 mm both rostral and caudal from the injured epicenter, CD1dKO mice had a significantly greater number of viable motor neurons than WT mice post-SCI (Fig. 2F, 2G). Thus, a CD1d deficiency confers protection to spinal cord neurons surrounding the SCI impact core. These results suggest that CD1dKO mice can be used as an animal model to study secondary injury progression in the traumatic penumbra containing viable tissue around the irreversibly damaged SCI core. Additionally, our results highlight rationales to study penumbral pathophysiology, which may lead to the development of novel therapeutic strategies to salvage penumbral spinal cord tissue following a SCI.

### A CD1d deficiency enhanced locomotor functional recovery following a SCI

Because we found that CD1dKO mouse spinal cords had less tissue damage and neuronal cell death after a SCI, we investigated whether they also had a better functional recovery than WT mice post-SCI. Thus, both WT and CD1dKO mice were subjected to a T10 moderate contusive SCI as described above (30). After the SCI, the mice were subjected to the Basso Mouse Scale (BMS) (31), Grid-Walking (32), and Rotarod tests (33) for evaluating locomotion, sensorimotor, and motor-coordination functions (Fig. 3A, 3B), respectively. On days 1 and 3 post-SCI, both WT and CD1dKO mice had similarly low BMS scores with complete paralysis of their hind limbs (Fig. 3C, left panel). However, on day 7 post-SCI, CD1dKO mice started to show significantly improved hind limb functional recovery compared to WT mice (Fig. 3C, left panel). This striking difference was also evident in videos showing the relative mobility of WT (Movie S1) and CD1dKO mice (Movie S2) 3 weeks post-SCI. This broad and important difference in the level of functional recovery improvement continued for over 4 weeks of BMS examination (Fig. 3C, left panel). Thus, CD1dKO mice exhibit a better locomotor functional recovery from a SCI than WT mice.

**Fig. 3.**
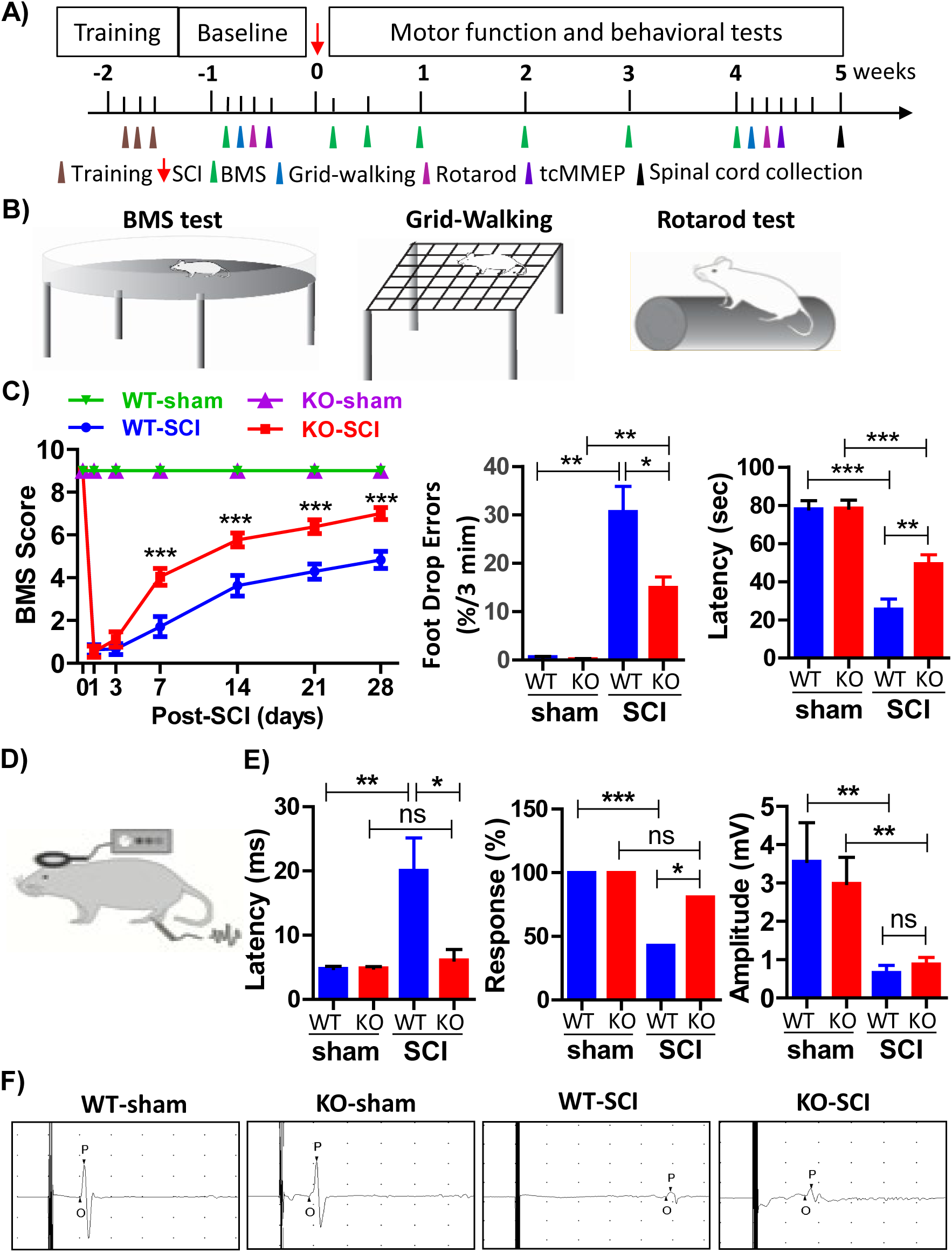
CD1dKO mice exhibited improved functional recovery following a SCI. **A)** and **B)** Experimental design and timepoints post-SCI for the BMS, Grid-Walking and Rotarod tests to evaluate locomotion deficits, sensorimotor deficits, and motor coordination. **C)** Pooled data showing BMS scores (left panel), foot drop errors (middle panel) and latency (right panel, 30 rpm) from WT and CD1dKO mice on day 28 post-SCI. **D)** tcMMEP to assess the functional integrity of the spinal cord’s descending axonal pathways following a SCI. **E)** Pooled data of tcMMEP showing the evoked potentials (left panel), total response rate (middle panel), and potential amplitude (right panel) from WT and CD1dKO mice on day 28 post-SCI. **F)** Representative recordings of tcMMEP showing the potential amplitude, latency and response rate in CD1dKO or WT mice on day 28 post-SCI or sham injury. Statistical analyses: two-way ANOVA (BMS), χ^2^ test (response), or a one-way ANOVA for all other groups. \**p*<0.05; \*\**p*<0.01; \*\*\**p*<0.001; ns, not significant. WT-SCI, n=12; CD1dKO-SCI, n=13; WT-sham, n=6; CD1dKO-sham, n=6.

The Grid-Walking test showed that both WT and CD1dKO mice post-SCI had an increased number of foot drop errors than those with a sham injury (Fig. 3C, middle panel). Compared to WT mice with a foot drop error rate of 26.7%, CD1dKO mice had a significantly lower foot drop error rate (15.0%) (Fig. 3C, middle panel), which constituted a 44% improvement over WT mice (*p*<0.01). Therefore, CD1dKO mice have a much greater recovery of sensory-motor coordination than WT mice after a SCI.

The Rotarod test showed that both WT and CD1dKO mice had a lower latency rate post-SCI than sham-injured mice over the 28-days of analysis (Fig. 3C, right panel). When compared to WT, CD1dKO mice had a significantly longer latency (Fig. 3C, right panel), suggesting CD1dKO mice have better motor coordination recovery post-SCI.

Transcranial magnetic motor-evoked potentials (tcMMEP) were used to assess the functional integrity of the spinal cord’s descending axonal pathways in both WT and CD1dKO mice following a SCI (Fig. 3D). WT mice exhibited a prolonged latency of the evoked potential (Fig. 3E, left panel), a lower total response rate (Fig. 3E, middle panel), and a reduced potential amplitude (Fig. 3E, right panel) post-SCI when compared to sham-injured WT mice. In marked contrast, injured CD1dKO mice only exhibited a reduction in potential amplitude, with no significant differences observed in either latency or response rate compared to sham-injured (Fig. 3E, 3F). Thus, of the three indicators of tcMMEP, spinal cord-injured CD1dKO mice only have a reduced potential amplitude, whereas WT mice exhibit deficits in all three post-SCI. This suggests that a CD1d-deficient environment *in vivo* preserves the descending axonal electrophysiological pathways in mice after a SCI.

Taken together, compared to WT, CD1dKO mice exhibit a remarkable improvement in locomotor functional recovery, sensory-motor coordination and balance, grip strength, hind limb motor coordination and functional integrity of the spinal cord post-SCI, indicating that CD1d negatively affects the functional outcomes of a SCI.

### Microglia/M*ɸ* of CD1dKO mice were less activated and had reduced proliferation in the injured spinal cord

Following a traumatic SCI, microglia/M*ɸ* undergo activation, proliferation, and changes in gene expression and morphology, exerting either detrimental or beneficial effects (34, 35). Given that spinal cord microglia/M*ɸ* constitutively expressed high levels of CD1d and this expression was further upregulated by a SCI, we hypothesized that CD1d had an impact on microglial/M*ɸ* responses to a SCI. To test this hypothesis, we performed IHC to analyze and compare the activation and proliferation of microglia/M*ɸ* by quantifying Iba1 expression and *in vivo* incorporation of 5′-bromo-2′-deoxyuridine (BrdU) in injured spinal cord tissues collected from WT and CD1dKO mice on day 7 post-SCI. In both WT and CD1dKO mice, Iba1 expression intensity and Iba1^+^ cell numbers were dramatically greater than those in sham-injured mice (Fig. 4A, 4B), indicating that a SCI triggers activation and proliferation of microglia/M*ɸ*. Interestingly, injured CD1dKO mice had fewer numbers of Iba1^+^ microglia/macrophages in the injured epicenter and 0.5 - 1.0 mm rostral and caudal from the injury site when compared to WT mice (Fig. 4A, 4B). This suggests that CD1d signaling promotes microglial/M*ɸ* proliferation. Similarly, when compared to injured WT mice, CD1dKO mice post-SCI had lower numbers of BrdU^+^ or BrdU^+^Iba1^+^ microglia/M*ɸ* in the injured epicenter as well as 1.0 mm both rostral and caudal from the injury site (Fig. 4A, 4C, 4D). To confirm these results, we also used single-cell RNA sequencing (scRNA-seq) to analyze and compare transcriptomes of spinal cord cells from WT and CD1dKO mice with or without a SCI. As shown in Fig. S2, microglia were highly activated in the spinal cords of WT mice following a SCI compared to sham-injured WT mice. In WT mice, injured CD1dKO mice expressed much lower levels of genes involved in microglia activation (Fig. S2B), confirming that CD1d is associated with microglia activation as revealed in our IHC analysis. Thus, CD1d-dependent signaling likely promotes the activation and proliferation of microglia/M*ɸ* in the spinal cord post-SCI. The lack of CD1d would then reduce activation and proliferation of microglia/M*ɸ*, which may be responsible for the neuroprotection that was observed in injured CD1dKO mice.

**Fig. 4.**
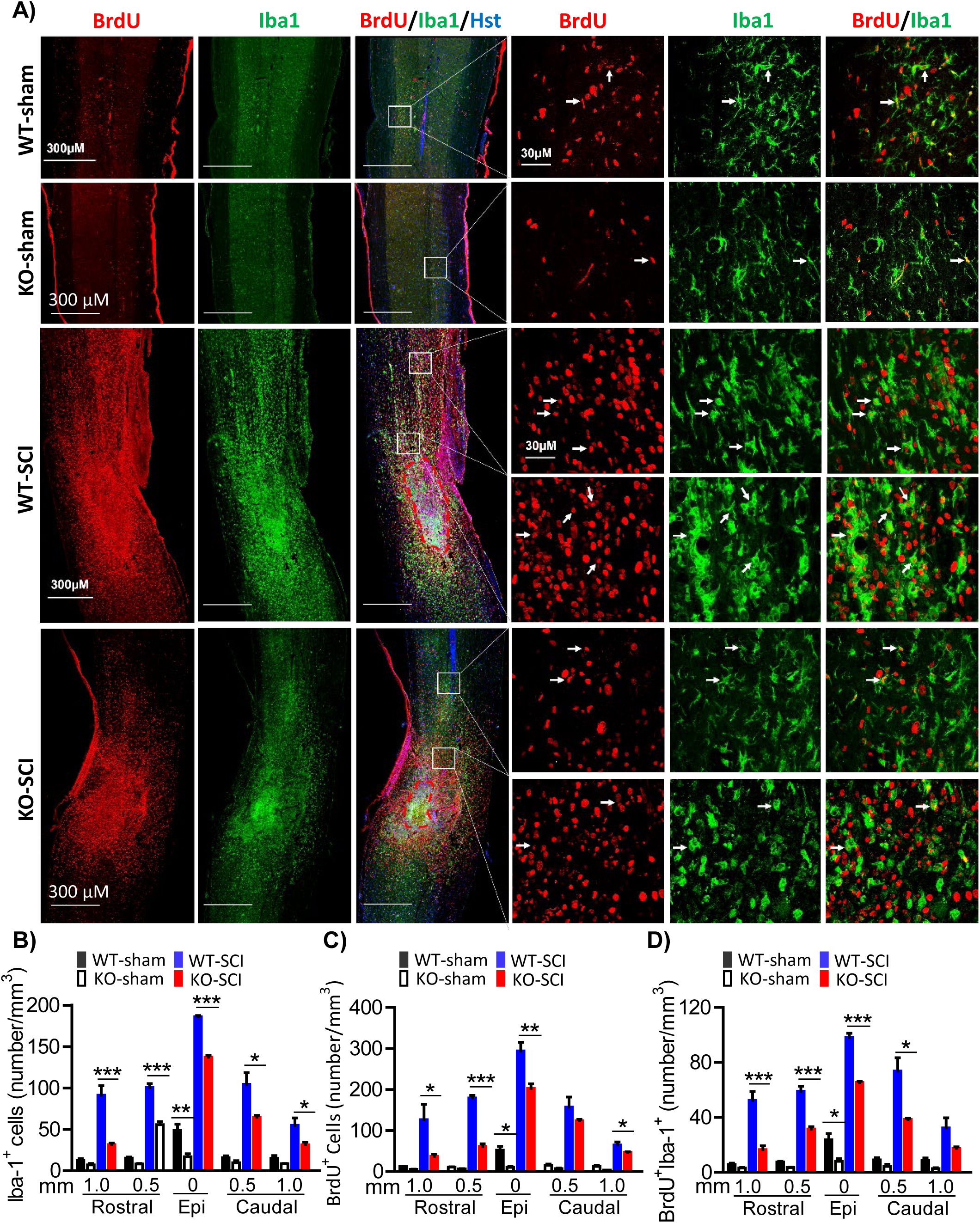
Effect of CD1d on SCI-triggered activation and proliferation of spinal cord microglia/macrophages *in vivo*. **A)** Injured spinal cord tissues from WT and CD1dKO mice, including the injured epicenter and 0.5-1.0 mm both rostral and caudal away on day 7 post-SCI. **B)**, **C)**, and **D)** numbers/mm^3^ of Iba1^+^, BrdU^+^ cells, and BrdU^+^Iba1^+^, respectively, in the injured epicenter and 0.5-1.0 mm rostral and caudal away. Student’s t-test was used for statistical analysis. \**p*<0.05, \*\**p*<0.01, \*\*\**p*<0.001. n=3/group. Epi, Epicenter. Hst, Hoechst dye (blue) for DNA staining.

### A CD1d deficiency promoted the polarization of macrophages into those involved in tissue repair and the anti-inflammatory response in the spinal cord post-SCI

A SCI triggers sterile neuroimmune/neuroinflammatory responses dominated by tissue-resident microglia and monocytes/macrophages (M/M*ɸ*)(10). These neuroimmune/neuroinflammatory responses can have both beneficial and detrimental effects (36–38). Here, we analyzed and compared M/M*ɸ*, as well as other types of immune cells in WT and CD1dKO spinal cords following a SCI. CD1dKO and WT mice post-SCI had a comparable percentage of neutrophils, B cells, total T cells, CD4 T cells, and CD8 T cells in the injured spinal cord (Fig. S1). Notably, the frequency of NKT cells (CD45^+^CD3^+^NK1.1^+^ T cells) was extremely low or negligible in spinal cord tissues (<0.17%) from WT mice with or without a SCI, but high (∼35%) in their livers (Fig. S1), suggesting that NKT cells act as key participants in liver physiology and pathology (39), but not in the spinal cord. In sharp contrast, the frequencies of spinal cord M/M*ɸ*, including inflammatory (CD45^+^CD11b^+^Ly-6G^-^Ly-6C^High^) and tissue-repairing (CD45^+^CD11b^+^Ly-6G^-^Ly-6C^Low^F4/80^High^), were dramatically affected by a SCI in both CD1dKO and WT mice. As shown in Fig. 5A and 5B, the frequencies of inflammatory and tissue-repairing M/M*ɸ* displayed opposite temporal patterns in injured spinal cord tissues from both WT and CD1d1 KO mice over the 28-day analyses post-SCI; specifically, the frequencies of inflammatory M/M*ɸ* were high in the initial acute phase of SCI and then gradually declined, whereas the frequencies of tissue-repairing M/M*ɸ* were low in the early phase of SCI and then increased over time (Fig. 5B, left panel). Interestingly, day 7 post-SCI is a pivotal turning point where the frequencies (%) of tissue-repairing M/M*ɸ* are significantly higher in injured spinal cord tissues from CD1d1 KO vs WT mice (Fig. 5C). In contrast, the frequencies of both inflammatory and tissue-repairing M/M*ɸ* in livers from WT and CD1dKO mice remained unchanged over the 28-day period post-SCI (Figure 5A, 5B right panel). These results suggest that M/M*ɸ* undergo differentiation and functional changes in response to the local spinal cord tissue-specific environment rather than systemic factors. Concurrently, day 7 post-SCI (post-acute phase) was also a turning point, when spinal cord lesions were significantly reduced while motor functions were markedly improved in CD1d1KO when compared to WT mice (Fig. 3). Thus, in CD1dKO mice, tissue-repairing M/M*ɸ* appear to be locally derived and substantially contribute to tissue repair and wound healing of the injured spinal cord following a SCI.

**Fig. 5.**
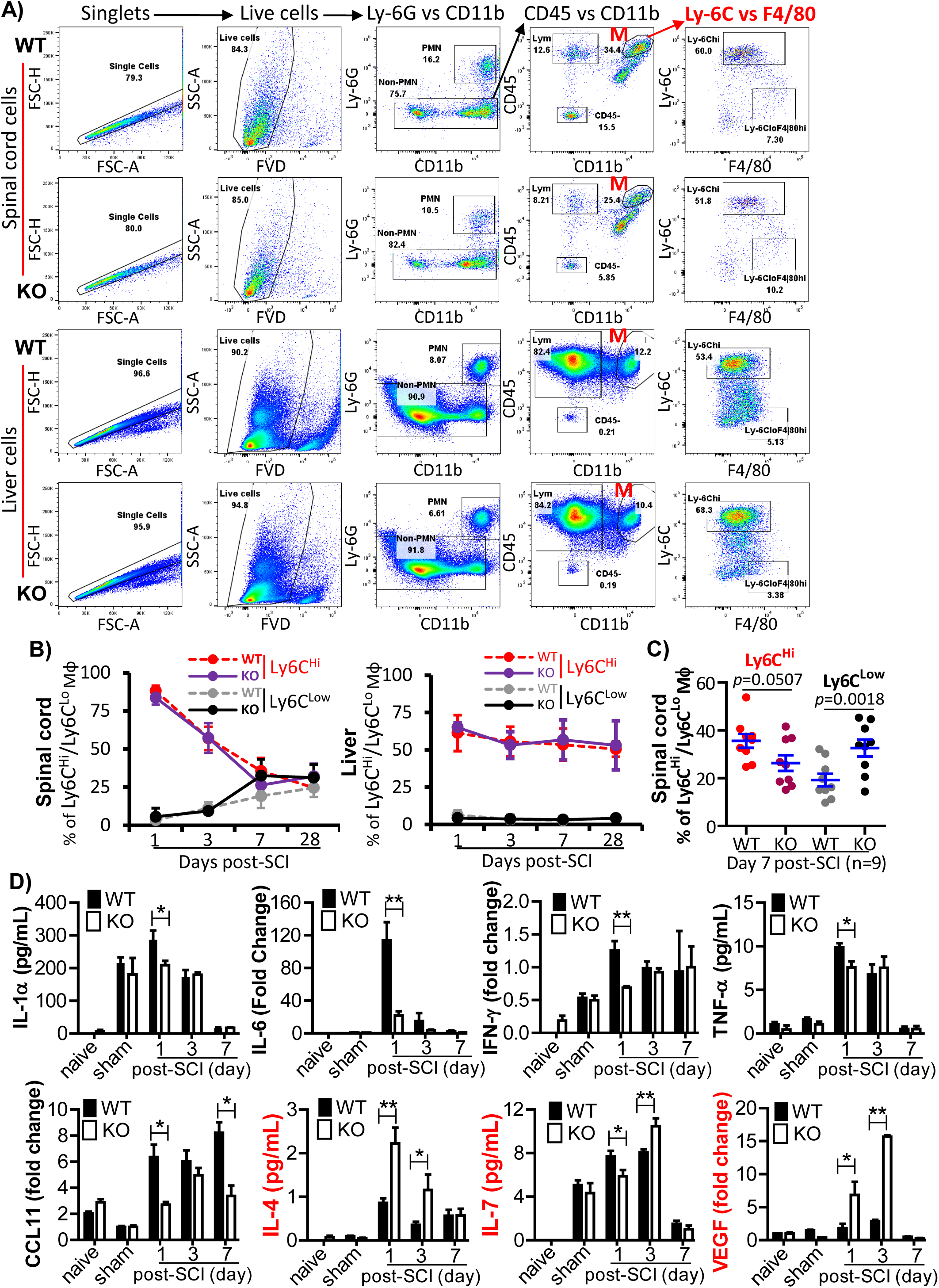
Effects of CD1d expression on the profiles of immune cells and pro-inflammatory/anti-inflammatory cytokines in the injured spinal cord post-SCI. At the indicated timepoints post-SCI, whole liver and 2 mm of spinal cord tissue including the injured epicenter and 1 mm rostral and caudal away were harvested. Single-cell suspensions were stained with Zombie Violet™ FVD to exclude FVD-positive dead cells, anti-CD16/CD32 mAbs for blocking FcR binding and fluorochrome-conjugated mAbs against mouse CD45, CD3, CD4, CD8, CD11b, Ly-6G, Ly-6C, F4/80, B220, and NK1.1. Tissue-infiltrating leukocytes (LYM) were defined as CD45^High^ cells. Within the CD45^High^ population, polymorphonuclear neutrophils (PMNs) were identified by Ly-6G expression, T cells as CD45^High^CD3^+^ cells, CD4 T cells as CD45^+^CD3^+^CD4^+^, CD8 T cells as CD45^+^CD3^+^CD8^+^, B cells as CD45^+^B220^+^, NKT cells (CD45^+^CD3^+^NK1.1^+^) and monocytes/macrophages (M/M*ɸ*) as CD45^+^CD11b^+^Ly-6G^−^. M/M*ɸ* were further divided into inflammatory Ly-6C^High^ and tissue-repairing Ly-6C^Low^F4/80^High^ subsets. Gating strategy for flow cytometric analysis of immune cells in the spinal cord (upper panel) and liver (lower panel) of WT vs CD1dKO mice post-SCI. The gated M/Mϕ population (M) showing the % of Ly-6C^High^ versus Ly-6C^Low^F4/80^High^ M/Mϕ in the spinal cord and liver. **B)** Pooled data of Ly-6C^High^ versus Ly-6C^Low^F4/80^High^ M/Mϕ in the spinal cord (left panel) and liver (middle panel) on day 1 (n=4/group), 3 (n=4/group), 7 (n=9/group) and 28 (n=4/group) post-SCI. Pooled data showing Ly-6C^High^ versus Ly-6C^Low^F4/80^High^ M/Mϕ in the spinal cords of WT and CD1dKO mice on day 7 post-SCI (n=9/group, right panel). **C)** Pooled data from the multiplex immunoassay showing the tissue levels of 8 proinflammatory/anti-inflammatory mediators that were significantly different between groups of WT and CD1dKO mice. Student’s t-test was used for statistical analysis. \**p*<0.05, \*\**p*<0.01. n = ≥4 per group.

To better understand the local tissue microenvironment that might determine the effects of CD1d on neuroimmune responses and the polarization of tissue-repairing M/M*ɸ* in response to a SCI, we simultaneously quantified spinal cord tissue levels of 32 cytokines, chemokines and growth factors using a multiplex immunoassay. After an extensive perfusion of the mice with PBS to remove intravascular cells, we harvested a 1-cm section of spinal cord (including the SCI site) for extracting proteins that were used for the multiplex immunoassay. Compared to naïve or sham conditions, an SCI caused significantly altered levels of eight of these analytes (Fig. 5D). Among these analytes, five proinflammatory cytokines/chemokines including IL-1α, IL-6, IFN-γ, TNF-α, and CCL11 (also known as Eotaxin) were significantly reduced in injured spinal cord tissues from CD1dKO when compared to WT mice (Fig. 5D); this mainly occurred in the early acute phase of SCI (day 1 post-SCI, Fig. 5D). These results are in accordance with the idea that CD1d signaling affects the inflammatory M/M*ɸ* response in the early phase of a SCI. Thus, the reduced levels of inflammatory cytokines/chemokines observed in CD1dKO vs WT mouse injured spinal cords are due to the CD1d-deficient environment limiting the development of inflammatory M/M*ɸ* in the spinal cord during the early phase of a SCI. Strikingly and of further important note, injured spinal cord tissue from CD1dKO mice had higher levels of IL-4, IL-7 and VEGF than that in WT mice (Fig. 5D). Moreover, the elevated levels of these analytes were maintained from the acute to subacute phase (day 1 - day 3) following a SCI (Fig. 5D), in which tissue-repairing M/M*ɸ* started to become the dominant subpopulation of M/M*ɸ* overall (Fig. 5A, 5B, 5C). IL-4 is a well-known anti-inflammatory cytokine and also orchestrates the differentiation of inflammatory M/M*ɸ* into tissue-repairing M/M*ɸ* (8, 40). IL-7 directly promotes neuronal survival and neuronal progenitor cell differentiation (41). VEGF is one of the most important proangiogenic mediators that promote cellular reactions throughout all phases of the wound healing process (42). Thus, the elevated IL-4, IL-7, and VEGF levels in CD1dKO spinal cord tissues confer anti-inflammatory and tissue-repairing benefits.

### Spinal cord neuropathology post-injury was not via the conventional CD1d/NKT cell axis

CD1d not only presents lipid antigens to NKT cells but can also exert intrinsic signaling for regulating immune and inflammatory responses (24–27). As we found that the frequency of NKT cells was extremely low or negligible in spinal cord tissues from WT mice, whether subjected to a SCI or not, we hypothesized that CD1d might contribute to the observed spinal cord neuropathology post-injury in a manner independent of the conventional CD1d/NKT cell axis. To test this hypothesis, we studied the effects on SCI outcomes of α-galactosylceramide (α-GalCer), a glycolipid that when presented by CD1d strongly activates Type I NKT cells *in vivo* (43). WT mice received α-GalCer or vehicle (PBS) via *i.p.* injection at 10 min after injury, every day of the first week and twice a week from the 2^nd^ to 6^th^ week post-SCI or sham injury (Fig. 6A). The mice were then subjected to the motor and behavioral tests described above. As shown in Fig. 6B, no significant differences in the BMS, Grid-Walking, and Rotarod tests were found between α-GalCer and PBS treatment groups at any of timepoints tested (Fig. 6B). Thus, α-GalCer-triggered activation of Type I NKT cells *in vivo* does not exhibit a significant impact on the motor and behavioral deficits that were observed in WT mice post-SCI.

**Fig. 6.**
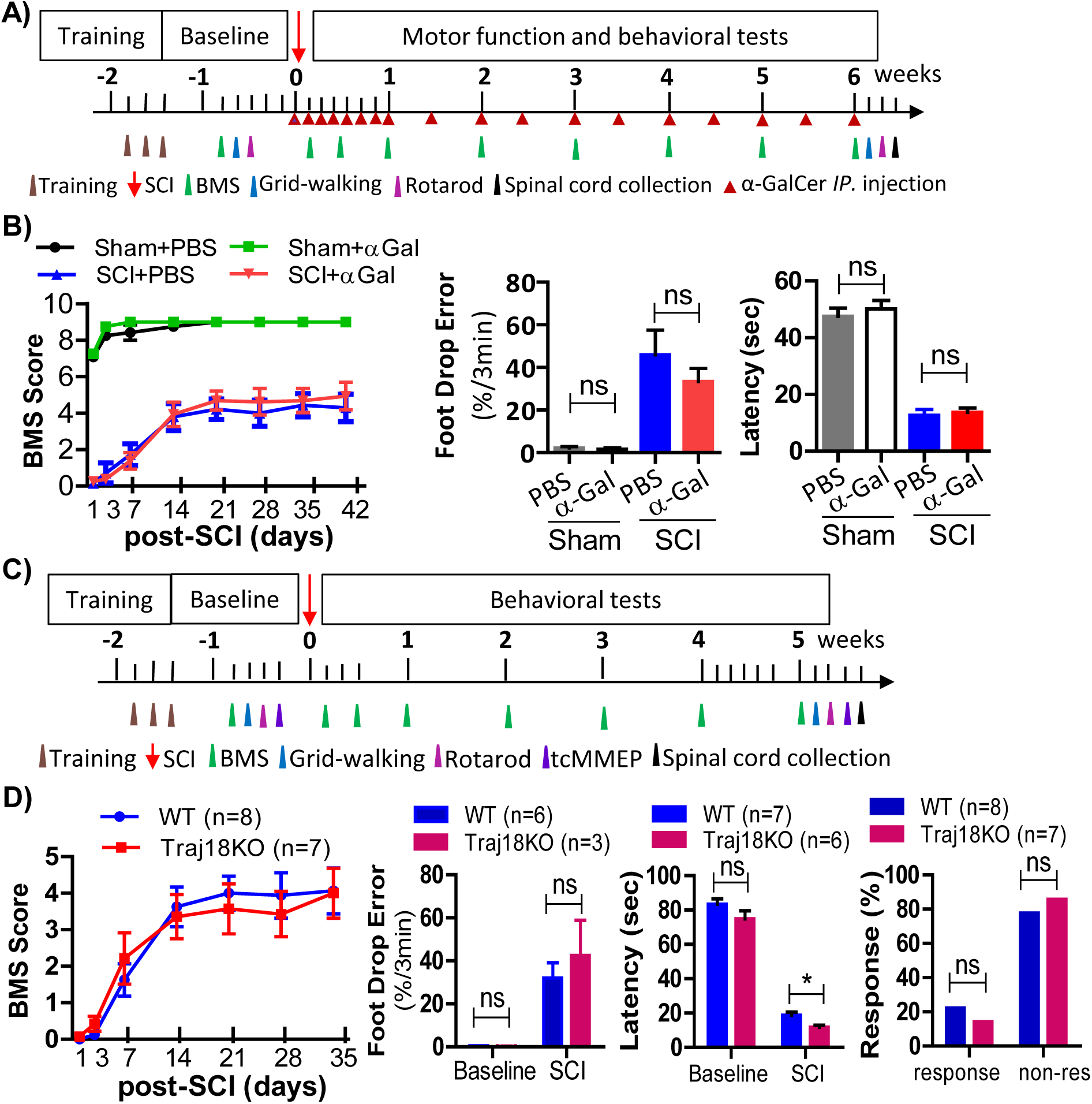
CD1d-dependent spinal cord neuropathology after a SCI was independent of the conventional CD1d/NKT cell axis. **A)** Layout and timeline for pre-injury training, SCI, α-GalCer administration via *ip.* injection, and motor and behavioral assessments of WT mice. **B)** Pooled data of BMS (left panel), Grid-Walking (middle panel), and Rotarod tests (right panel). **C)** The layout and timeline for pre-injury training, SCI and, motor and behavioral assessments of WT and Traj18KO (Type I NKT cell-deficient) mice. **D)** Pooled data of BMS (left panel), Grid-Walking (middle panel), and Rotarod tests (right panel). A two-way ANOVA was used for BMS data analysis. Student’s *t*-test was used for analyzing Grid-Walking and Rotarod test results. \**p*<0.05. WT-SCI, n=6; Traj18KO-SCI, n=7; sham + PBS, n=6; sham + α-Gal, n=6; SCI + PBS, n=7; SCI + α-Gal, n=8. α-Gal: α-GalCer; non-res: non-response.

To further clarify the role of the CD1d/NKT cell axis in the pathogenesis of SCI, we also studied SCI outcomes in C57BL/6 *Traj18* knockout (Traj18KO) mice that selectively lack Type I NKT cells due to a genetic deletion in the *Traj18* gene encoding the J region of their NKT cell receptor α chain (44). WT and Traj18KO mice were subjected to a T10 contusive SCI followed by motor and behavioral tests as described above (Fig. 6C). Additionally, a tcMMEP test was conducted on the right and left hind limbs of WT and Traj18KO mice before and after a SCI. No significant differences in the BMS, Grid-Walking, and tcMMEP assessments were found between the groups of WT or Traj18KO mice at any time point post-SCI (Fig. 6D), although Traj18KO mice had a shorter latency than WT mice on the Rotarod 5 weeks post-SCI (Fig. 6D). Thus, almost all motor and behavioral measurements are comparable between WT and Traj18KO mice post-SCI, suggesting that the conventional CD1d/NKT cell axis is unlikely to be involved in the CD1d-mediated spinal cord neuropathology observed following a SCI.

### Administration of an anti-mouse CD1d mAb ameliorated CD1d-mediated spinal cord neuropathology following a SCI

Lipids are an essential structural and functional components of all cell types and tissues in the human body (12). Particularly, the CNS has a rich lipid composition with about 75% of all mammalian lipid species that are exclusively present in neural tissues (14), demonstrating the unique requirements of lipids for neural cell functions (15, 16). In SCI, damaged neural cells and tissues release lipids such as glycolipids that represent a dominant class of lipids in the myelin bilayer of neural cells (17). Released glycolipids such as CD1d ligands may directly interact with CD1d on the surface of CD1d^+^ spinal cord cells (microglia/M*ɸ*, ODCs, and ECs) to trigger CD1d intrinsic signaling in a manner independent of the canonical CD1d/NKT cell axis. To clarify whether blockade of CD1d-lipid interaction could confer a neuroprotective effect, we injected an anti-mouse CD1d mAb (1H6, IgG2b)(45) or isotype (IgG2b) control i.p. to WT mice before and after a SCI as illustrated in Fig. 7A. These mice were subjected to the BMS, Grid-Walking, and Rotarod tests to evaluate the effects of the anti-mouse CD1d mAb on motor and behavioral outcomes post-SCI (Fig. 7A, 7B). As shown in Fig. 7B, treatment with the anti-CD1d mAb significantly increased BMS scores (Fig. 7B, left panel), decreased the hind limb foot drop error rate (Fig. 7B, middle panel), and increased latency on the Rotorod at 6 weeks post-injury (Fig. 7B, right panel). Injured spinal cord tissues were collected on day 45 post-SCI for IHC analysis of lesion area size and the amount of SWM (Fig. 7C, 7D). Treatment with the anti-CD1d mAb significantly reduced the lesion area size and resulted in a greater amount of SWM compared to injured mice receiving the isotype control mAb (Fig. 7C, 7D). These results demonstrate that treatment with this anti-CD1d mAb has comparable neuroprotective effects as those observed in CD1dKO mice, suggesting that a CD1d blockade can ameliorate CD1d-mediated spinal cord neuropathology and improve outcomes following a SCI.

**Fig. 7.**
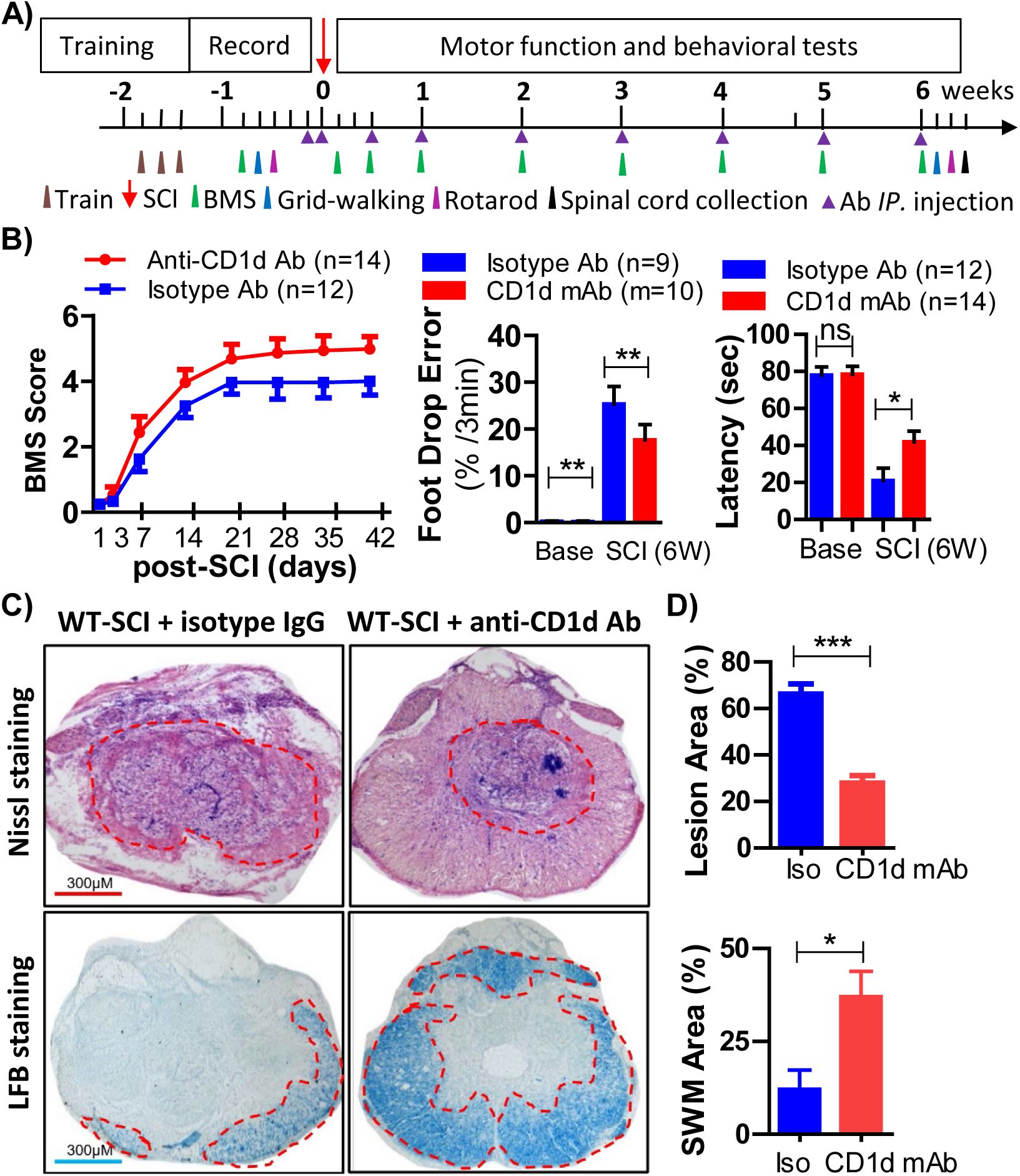
Blockade of CD1d signaling ameliorated CD1d-mediated spinal cord neuropathology of SCI. **A)** Layout and timeline for pre-injury training, SCI, anti-mouse CD1d mAb administration via *i*.*p.* injection, spinal cord tissue collection, and motor and behavioral assessments of WT and CD1dKO mice. Anti-mouse CD1d (1H6) or an isotype control mAb was administered *i*.*p*. (50 µg/mouse) into WT mice at various times post-SCI. **B)** Pooled data of BMS (left panel), Grid-Walking (middle panel) and Rotarod tests (right panel). **C)** Representative images of Nissl (upper panel) and LFB staining (lower panel) on histological changes of spinal cord tissue lesion size and spared white matter (SWM). **D)** Pooled data showing spinal cord tissue lesion size (upper panel) and spared white matter volume (lower panel) in SCI mice treated with the anti-CD1d mAb or isotype control mAb. Two-way ANOVA was used for analyzing the BMS data. Student’s t-test was used for analyzing foot drop errors, latency, lesion area and amount of spared white matter (SWM). \**p*<0.05, \*\**p*<0.01, \*\*\**p*<0.001. CD1d mAb + WT-SCI, n=14; isotype mAb + WT-SCI, n=12.

## Discussion

This report provides compelling evidence that CD1d, an MHC class I-like molecule mostly known for presenting lipid antigens to NKT cells, plays an important role in the neuropathology that occurs following a SCI. We found that CD1d was constitutively expressed on microglia/M*ɸ*, ODCs, and ECs, but not on either neurons or astrocytes, in the spinal cord. Moreover, this CD1d expression was upregulated after a SCI. In the spinal cord post-injury, as compared to WT, CD1dKO mice exhibited less tissue damage, reduced activation and proliferation of microglia/M*ɸ*, and lower levels of proinflammatory cytokines, but higher levels of anti-inflammatory cytokines and increased frequencies of tissue-repairing M*ɸ*. Consequently, CD1dKO mice had substantially improved post-SCI functional recovery vs WT mice. The contribution of CD1d in spinal cord neuropathology after a SCI occurs in a manner independent of the conventional CD1d/NKT cell axis because: (***1***) the frequency of NKT cells was extremely low or negligible (∼0.17%) in spinal cord tissues of WT mice with or without a SCI, ***(2)*** Spinal cord-injured WT mice receiving α-GalCer, a glycolipid that when presented by CD1d strongly activates NKT cells (46), were no different in their motor function and behavioral assessment scores when compared to the control group, suggesting *in vivo* activation of NKT cells by α-GalCer had no significant impact on the functional and behavioral deficits observed in WT mice post-SCI, and (***3***) there were no differences in motor function and behavioral assessments after a SCI between WT mice and C57BL/6 Traj18KO mice, with the latter selectively lacking Type I NKT cells (44). These results suggest that the CD1d-associated neuropathology following SCI is mediated by CD1d intrinsic signaling or unknown mechanisms rather than the canonical CD1d/NKT cell axis.

As an MHC class I-like glycoprotein that is primarily expressed on antigen-presenting cells (APCs), CD1d consists of two polypeptide chains (α and β2-microglobulin or β2M). CD1d interacts with T cell receptor (TCR) on the surface of NKT cells to present lipid antigens. In contrast to the classical MHC class I molecules that are highly polymorphic, CD1d molecules exhibit limited sequence diversity (47). CD1d has a short cytoplasmic tail containing a putative tyrosine-dependent signal motif YXXZ (where Y signifies tyrosine, X is any amino acid, and Z is a hydrophobic amino acid) (25, 48). Crosslinking human CD1d on the surface of myeloid cells induces tyrosine phosphorylation and cytokine production (25). These crosslinking effects are abolished when the CD1d cytoplasmic tail is replaced with a CD1a cytoplasmic domain (25), suggesting that CD1d exerts intrinsic signaling via its YXXZ motif in the cytoplasmic tail. A recent study has also revealed that CD1d intrinsic signaling in macrophages controls NLRP3 inflammasome expression during inflammation in an animal model of dextran sodium sulfate (DSS)-induced colitis (24). Mechanistically, CD1d engagement with the natural glycosphingolipid ligand iGb3 induces intracellular Ser^330^ dephosphorylation, which reduces AKT-dependent STAT1 phosphorylation and subsequent NF-*κ*B activation, eventually leading to the transcriptional down-regulation of NLRP3 inflammasome expression in macrophages in DSS-induced colitis (24). Therefore, CD1d not only binds and presents lipid antigens for recognition by NKT cells but also exerts intrinsic signaling to regulate inflammatory and immune responses (24, 25). Given our findings that CD1d-associated neuropathology following a SCI was independent of the canonical CD1d/NKT cell axis, CD1d intrinsic signaling was likely triggered. In this regard, CD1dKO and WT mice post-SCI could be used to investigate the pathogenic role of CD1d intrinsic signaling in SCI-triggered sterile neuroimmune and neuroinflammatory responses in the spinal cord, which warrants further studies.

In mice, there are two broad subsets of M/M*ɸ* known as inflammatory (CD45^+^CD11b^+^Ly-6G^-^Ly-6C^High^) and tissue-repairing (CD45^+^CD11b^+^Ly-6G^-^Ly-6C^Low^F4/80^High^) (49). During inflammation or tissue injury, M/M*ɸ* responses are characterized by the accumulation of inflammatory M/M*ɸ* in the first phase to clear cellular debris, followed by the conversion to tissue-repairing M/M*ɸ* in the reparative phase that contributes to tissue repair and wound healing (9). The phenotypic switch from inflammatory to tissue-repairing M/M*ɸ* is mainly driven by IL-4 at the site of inflammation or injury (8, 9), which is more efficient than systemic differentiation factors (50). Inflammatory M/M*ɸ* express higher levels of proinflammatory cytokines such as IL-1, IL-6, IL-8, and TNF-α during the early phase of inflammation or tissue injury (6, 7), whereas tissue-repairing M/M*ɸ* produce growth factors such as VEGF and anti-inflammatory mediators (e.g., IL-10 and TGF-β1) that foster inflammation resolution, tissue repair, and wound healing (8, 9). Strikingly, compared to WT mice, CD1dKO mice had a significantly higher frequency of tissue-repairing M/M*ɸ* in injured spinal cord tissues in the post-acute phase of SCI (Fig. 5). Indeed, the frequencies of inflammatory and tissue-repairing M/M*ɸ* displayed opposite temporal patterns in injured spinal cord tissues from both WT and CD1d1 KO mice over the 28-day course of SCI studied here (Fig. 5). Specifically, the frequencies of inflammatory M/M*ɸ* were high in the initial phase of SCI and then gradually declined, whereas the frequencies of tissue-repairing M/M*ɸ* were low in the early phase of SCI and then increased over time (Fig. 5). However, day 7 post-SCI was a turning point where the frequency of tissue-repairing M/M*ɸ* was significantly higher in injured spinal cord tissues from CD1dKO mice than those from WT (Fig. 5). In contrast, the frequencies of both inflammatory and tissue-repairing M/M*ɸ* in WT and CD1dKO mouse livers remained unchanged over the 28-days post-SCI (Fig. 5). These results suggest that M/M*ɸ* undergo differentiation and functional changes in response to local tissue-specific factors rather than from systemic signals. Concomitantly, day 7 post-SCI was also a turning point in CD1dKO vs. WT mice when inflammation and lesions in spinal cord tissue were significantly reduced (Fig. 2), whereas motor function was markedly improved (Fig. 3). Thus, tissue-repairing M/M*ɸ* appear to be derived by local factors in the spinal cord microenvironment and make a substantial contribution to tissue repair and wound healing of the injured spinal cord in CD1dKO mice following a SCI.

We also found that CD1dKO mice had higher levels of IL-4, IL-7, and VEGF in injured spinal cord tissues post-SCI than WT mice (Fig. 5). IL-4 is a well-known anti-inflammatory cytokine and also orchestrates the differentiation of inflammatory M/M*ɸ* into tissue-repairing M/M*ɸ* (8, 40). IL-7 promotes both neuronal survival and neuronal progenitor cell differentiation (41) and VEGF is one of the most important proangiogenic cytokines, promoting cellular reactions throughout all phases of the wound healing process (42). Thus, the elevated levels of IL-4, IL-7, and VEGF in CD1dKO injured spinal cord tissues induce the polarization of inflammatory M/M*ɸ* into tissue-repairing M/M*ɸ* and the consequent tissue-repairing benefits. In addition to the tissue milieu in which IL-4 amplifies tissue-repairing M/M*ɸ* polarization, Nr4a1 (nuclear receptor subfamily 4 group A member 1, also known as Nur77) is a transcription factor that not only plays a key role in the differentiation of tissue-repairing M/M*ɸ* from inflammatory M/M*ɸ,* but it is also essential for the survival of tissue-repairing M/M*ɸ* (51, 52). Our scRNA-seq analysis (Fig. S2B) revealed that Nr4a1 was highly expressed in M*ɸ*, especially in those from WT or CD1dKO mice following a SCI. In addition, VEGF-A was predominantly produced by M2 rather than M1 M*ɸ*. Thus, abrogation of CD1d signals likely upregulates SCI-induced gene expression of VEGF-A and Nr4a1, thereby increasing survival and differentiation of M2 macrophages (or tissue-repairing M/M*ɸ*) to promote tissue repair and regeneration. Of note, the levels of MCP-1 (also known as CCL2) in injured spinal cord tissues from WT and CD1dKO mice post-SCI were comparable (Table S1). Given that MCP-1 is a principal chemokine for recruiting Ly-6C^High^ M/Mϕ into inflammatory sites through its binding to its receptor CCR2 that is highly expressed on inflammatory Ly-6C^High^ M/M*ϕ* (53, 54), the similar levels of MCP-1 in injured spinal cord tissues from CD1dKO and WT mice explain the comparable frequencies of inflammatory M/Mϕ found in the acute phase of SCI in these groups.

CD1d ligation alone, in the absence of NKT cells, can rapidly trigger activation of the NF-κB pathway in CD1d^+^ cells such as human M/Mϕ and dendritic cells (DCs), and subsequently induces cytokine production and inflammasome activation (24, 25, 55). These findings imply that surface CD1d on M/Mϕ, ODCs, and ECs in the spinal cord can be directly engaged by lipids released from damaged tissues and cells post-SCI. In this regard, direct interaction of CD1d with lipids can be blocked. Indeed, we found that administration of anti-mouse CD1d blocking mAb via *i.p.* injection to WT mice before and after SCI ameliorated CD1d-mediated spinal cord neuropathology (Fig. 7). Treatment with this anti-CD1d mAb significantly increased BMS scores, decreased the hind limb foot drop error rate and increased latency on the Rotorod 6 weeks post-SCI (Fig. 7). IHC analysis also showed that treatment with the anti-mouse CD1d mAb significantly reduced the size of the lesion area and protected white matter (Fig. 7). In fact, treatment with our anti-mouse CD1d mAb had comparable neuroprotective effects as those observed in CD1dKO mice, thereby representing a potential immunotherapeutic strategy for SCI. That being said, although we were able to demonstrate that Type I NKT cells were not involved in the CD1d-dependent neuropathology post-SCI, we cannot rule out a potential role for Type II (or variant) NKT cells, as it has been shown previously that CD1d can present the CNS lipid sulfatide, which can activate at least one subpopulation of Type II NKT cells (20, 56–59). Regardless, for potential translational use of a CD1d-specific targeted therapy in SCI, further work is warranted to determine the optimal dose, optimal time window and side effects of administering the CD1d-specific mAb used in the current study.

Taken together, our results suggest that a SCI triggers a Type I NKT cell-independent, CD1d-dependent neuroimmune/neuroinflammatory response that recruits inflammatory M/M*ɸ* into inflamed spinal cord tissues during the early phase of SCI. In a CD1d-deficient environment, there would be reduced CD1d-intrinsic proinflammatory signaling, but an increase in IL-4 production that promotes the differentiation of inflammatory M/M*ɸ* into the tissue-repairing variety. Tissue-repairing M/M*ɸ* produce high levels of IL-4 and VEGF which counteracts neuroinflammation and improves tissue repairing, respectively. Therefore, our findings provide fundamental insight into the mechanisms underlying SCI-triggered sterile neuroimmune/neuro-inflammatory responses in the spinal cord. Targeting CD1d may represent a novel therapeutic strategy to reduce spinal cord damage and improve functional recovery after a SCI.

## Materials and Methods

### Animals

Eight-week-old C57BL/6J wildtype (WT) male and female mice, and Sprague Dawley (SD) adult and pregnant rats were purchased from Charles River (Wilmington, MA). CD1d knockout (CD1dKO) mice on the C57BL/6J background (30) were originally provided by Dr. Luc Van Kaer (Vanderbilt University). C57BL/6 Traj18KO mice (44) were purchased from The Jackson Laboratory (Bar Harbor, ME).

### Primary spinal cord cell cultures

The generation of primary spinal cord neurons from E15 rats was performed as published in our previous work (60) and were cultured in DMEM supplemented with 10% FBS, 5% horse serum and 2mM glutamine (GE Healthcare Life Sciences) on 1.2 cm poly-L-lysine-coasted microscope coverslips before fixation and staining for immunofluorescent (IF) microscopy staining.

Glial cells were isolated from postnatal day 1 (P1) rat spinal cords and cultured in DMEM containing 10% FBS and 2 mM glutamine on coated T75 flasks as previously described (61). The cells were placed on coverslips and fixed for IF microscopy as described above.

### Contusive spinal cord injury

A moderate contusive SCI in mice was produced at the 10^th^ thoracic vertebra (T10) by the Louisville Injury System Apparatus (LISA) impactor (62). A T10 laminectomy was performed on anesthetized mice and a SCI induced by the LISA impactor. Sham-injured control animals received the laminectomy without the SCI.

### *In vivo* BrdU labeling

WT and CD1dKO male and female mice were given BrdU in their drinking water (5 g/L) and regular food immediately after a SCI for one week as previously described (63). The animals were then sacrificed, and the spinal cords were collected. Cryostat longitudinal sections (25 µM) were then cut for the histological assessments described in the text.

### Cytokine assay

A 1-cm piece of spinal cord containing the injured epicenter from PBS-perfused mice was removed and placed individually into a tube which contained 200 µl of PBS (0.01 M). The spinal cords were homogenized thoroughly and centrifuged at 40,000 rpm at 4°C for 15 min. The supernatants were collected for the detection of cytokines using the Milliplex MAP Mouse Cytokine/Chemokine premixed 32 magnetic bead panel (Millipore, Burlington, MA) following the manufacturer’s instructions. The data were acquired on a Luminex platform and were analyzed by MILLIPLEX Analyst 5.1 software (Millipore, Burlington, MA).

### qPCR of spinal cord mRNA

Total RNA was isolated using an RNAeasy kit (Qiagen, Hilden, Germany) according to the manufacturer’s protocol. Random hexamers and reverse transcriptase were used for cDNA synthesis (Roche Applied Science Transcriptor High Fidelity cDNA Synthesis Kit). Predesigned TaqMan probes for murine CD1d (Mm00783541) and GAPDH (Mm99999915_g1) were purchased from Applied Biosystems (Waltham, MA). CD1d and GAPDH transcripts were quantified using a TaqMan PCR Master Mix and ABI Prism 7000 instrument (Applied Biosystems, Waltham, MA). The relative change in *cd1d1* gene expression between the sham and SCI groups was calculated using the ΔC_t_ method (62) and represented as the fold change, which was normalized to the sham group.

### Behavioral assessments

Four behavioral assessments, the Basso Mouse Scale (BMS)(31), Grid-Walking (32), Rotarod (33) and transcranial magnetic-motor evoked potential (tcMMEP)(64), were used to evaluate hind limb functional recovery. Prior to surgery, each mouse was trained with the open field (BMS), Grid-Walking, and Rotarod every day for 3 days. The baselines of the behavioral tests were recorded.

### Histological assessments

Mice were anesthetized and perfused with PBS and 4% paraformaldehyde (PFA). A 3 cm segment of the spinal cords which included the injury epicenter was dissected, removed, and placed in 4% PFA overnight. The next day, the spinal cord segments were transferred into a 30% sucrose solution in 1X PBS for one week (65). One centimeter of spinal cord around the injury epicenter was embedded with OCT embedding medium (Fisher HealthCare, Waltham, MA) and cut into 25 µm thick cross or longitudinal sections, which were mounted on frosted slides.

### Lesion area and spared white matter measurement

One set of sections/slides was stained with Cresyl Violet Eosin for measuring the lesion area, lesion volume and viable neurons; another set was stained with Luxol Fast Blue for the assessment of the spared white matter area and volume (66). The epicenter of each spinal cord was selected, the cord area and lesion area were outlined, and measured with an Olympus BX60 microscope (Olympus, Tokyo, Japan) using Neurolucida Neuron Tracing Software (MicroBrightField, Williston, VT). The lesion area was calculated, and the area of spared white matter (SWM) was also measured using the Neurolucida software. The SCI SWM areas were compared to WT and CD1dKO sham white matter areas; the SWM areas (presented as percent) were then calculated (66).

### 3-Dimensional lesion and SWM volume assessments

The epicenters of the lesions were set as a middle (zero) point and 0.3 mm intervals, from the rostral to caudal areas of the spinal cord. The lesion and SWM volumes were calculated following published methods (67). The SCI length was calculated from the location at which the first section showed a lesion, to the location that was the end of the lesion from the rostral to caudal sides. A 3-Dimensional model was reconstructed with Neurolucida System Software (68).

### Immunofluorescence

The coverslips of cells or slides with tissue sections were incubated in 0.3% Triton X-PBS (PBST) twice for 15 min each, then incubated with 10% goat serum (Cell Signaling Technology) in PBST for 1 h. The primary antibodies used were specific for CD1d (BioLegend, San Diego, CA), βIII-tubulin (Millipore, Burlington, MA), NeuN (Millipore, Burlington, MA), GFAP (Abcam, Cambridge, United Kingdom), Iba-1 (Abcam, Cambridge, United Kingdom), CD31 (ThermoFisher, Indianapolis, IN) and Olig2 (Millipore, Burlington, MA). The next day, the coverslips and slides were incubated with the secondary antibodies, Cy3 goat anti-rat IgG and Alexa Fluor488 anti-rabbit IgG (Jackson ImmunoResearch, West Grove, PA), respectively, for 1 h at room temperature. Then, 5 µg/ml Hoechst (Cell Signaling Technology, Danvers, MA) in PBS was applied. Fluoromount-G (SouthernBiotech, Homewood, AL) was used as the mounting medium. For double staining with BrdU, the tissue sections were first stained with anti-Iba-1 and secondary antibodies, and then for BrdU (Abcam, Cambridge, United Kingdom). The secondary antibody (Cy3 goat anti-rat IgG) was applied, the slides were washed with PBS, incubated with Hoechst and mounted with fluoromout-G (69). After staining, images were taken with an Olympus BX60 microscope, an Olympus BX61 confocal microscope (FLUOVIEW, FV1200, Japan) and analyzed with ImageJ software (NIH, Bethesda).

### *In vivo* treatment with CD1d-specific mAb

WT male and female mice were randomly divided into two groups: SCI that received an isotype control antibody (TW2.3)(70), kindly provided by Dr. J. Yewdell, NIH, and those receiving the anti-mouse CD1d-specific mAb, 1H6, that we generated (45). Each mouse received 50 μg of either 1H6 mAb or TW2.3 mAb via *i.p.* injection and the mAb administration timing was as is illustrated in Fig. 7A. All the mice received BMS, Grid-Walking and Rotarod training before the SCI. Timing of mAb injections (i.p.) were as indicated. All mice were sacrificed on day 45 following SCI and spinal cord samples were collected as described above. Serial cryostat cross sections were cut and collected on the slides as above.

### Preparation of single cell populations from mouse spinal cord and liver

Sham- and SCI-WT mice were anesthetized and perfused with 0.01 M PBS as described above. For each mouse, a 2-cm section of spinal cord containing the injured site was collected and processed into a single cell suspension. After centrifugation at 700 x g for 10 min, the cells were resuspended in 10 ml of 40% Percoll (GE Healthcare, Chicago, IL) and centrifuged again at 950 x g for 20 min at room temperature. The cell pellets were harvested and used for flow cytometric analysis.

To prepare liver mononuclear cells (LMNCs), after perfusion, the liver was collected and ground through 70 µm cell strainers. The cells were centrifuged at 700 x g for 10 min and then resuspended in 10 mL of 40% Percoll, where they were centrifuged again at 950 x g for 20 min at room temperature. The cell pellets were harvested and resuspended in 5 mL of red blood cell (RBC) lysis buffer (BioLegend, San Diego, CA) for 5 min at room temperature. After centrifugation at 700 x g for 10 min at 4°C, the LMNCs were used for flow cytometric analysis.

### Flow cytometry

Single-cell suspensions of spinal cord or LMNCs were subjected tp surface staining to determine the frequency and phenotype of immune cells in these tissues. Briefly, cells were first stained with the Zombie Violet™ Fixable Viability dye (FVD, BioLegend, San Diego, CA) to exclude FVD-positive dead cells and FcR blocking reagent (eBioscience, San Diego, CA) to block non-specific binding to mouse Fc receptor, followed by staining with fluorochrome-conjugated antibodies (Abs) against mouse CD45, CD11b, Ly-6G, Ly-6C, F4/80, B220, CD3, CD4, CD8, and NK1.1. All Abs were purchased from BioLegend (San Diego, CA). The data were analyzed using FlowJo v10 software (Tree Star, San Carlos, CA).

### α-GalCer treatment

WT mice received a T10 moderate contusive SCI with the LISA impactor. Five minutes after the SCI, every day the first week and twice a week from the 2^nd^ to 6^th^ week, 100 µl (200 µg/mouse) of α-GalCer (ENZO Life Sciences, Farmingdale, NY) was injected i.p. The vehicle control was 100 µl of 1X PBS, i.p. injection. The BMS, Grid-Walking, and Rotarod tests were performed on the indicated days after the SCI.

### Traj18KO mouse SCI study

C57BL/6 background Traj18KO (Type I NKT cell-deficient) mice and wildtype (WT) littermates received training and surgery as described above. The behavioral tests were performed as described above.

### Statistics

Statistical analyses were carried out using GraphPad Prism software (version 7.00, LaJolla, CA) or R studio, as indicated in the Figure Legends. A *p* value below 0.05 was considered significant.

## Acknowledgements

We thank Drs. Luc Van Kaer and Jonathan Yewdell for the CD1dKO mice and TW2.3 hybridoma, respectively, and Dr. Mark Kaplan for comments on the manuscript. We also acknowledge the Flow Cytometry Resource Facility (FCRF) and the Center for Medical Genomics (CMG) in the Indiana University School of Medicine for their outstanding technical support. The Indiana University Melvin and Bren Simon Comprehensive Cancer Center FCRF is funded in part by National Cancer Institute grant P30 CA082709 and the National Institute of Diabetes and Digestive and Kidney Diseases grant U54 DK106846. The FCRF is also supported in part by NIH instrumentation grant 1S10D012270. This work was supported by NIH grants R01 NS103481 and R01 NS100531 (XMX), R01 NS111776 (WW), grants from the Indiana Traumatic Spinal Cord and Brain Injury Research Grant Program (RRB and WW), and NIH grant from UH2/UH3 AA026218 (QY). We dedicate this paper to our colleague and friend, Dr. Xiao-Ming Xu.

## Author Contributions

XW, QY, XMX, and RRB designed research; XW, JL, WL, MFK, HD, JT, RP, DJT, WW, AY, JG, FS, and XG performed research; XW, JL, WL, MFK, HD, JT, RP, DJT, WW, AY, JG, CHY, XG, FS, and QY analyzed data; XW, QY, XMX, and RRB wrote the paper.

## Classification

This article is a PNAS Direct Submission

