## supplementary figures 1 & 2 and supplementary Table 1 for "CD1d-dependent neuroinflammation impairs tissue repair and functional recovery following a spinal cord injury"

### Supplementary Figure 1

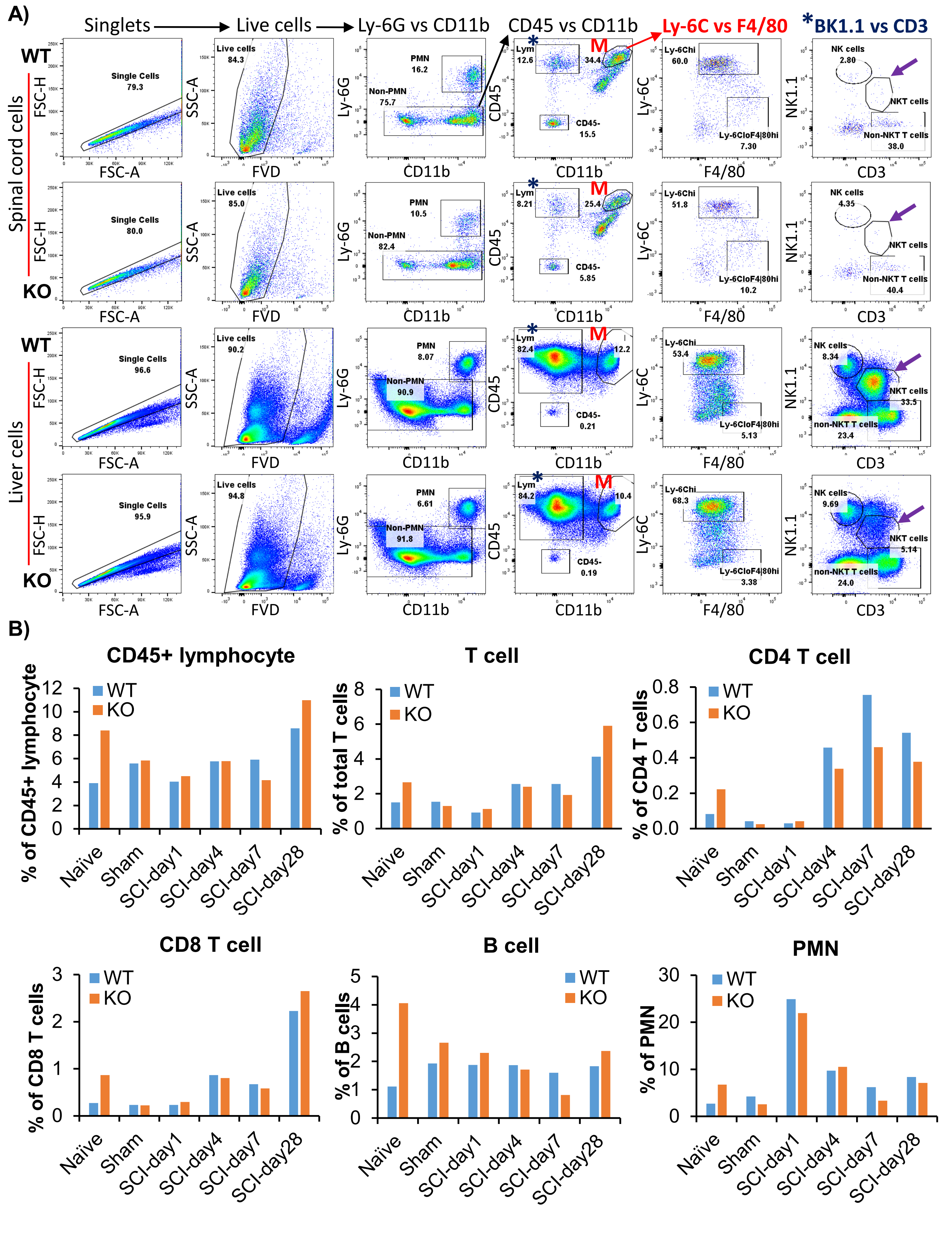

**Figure S1. Effects of a CD1d deficiency on immune cell profiles in the injured spinal cord post-SCI.** After a SCI, mice were perfused with heparinized PBS and then whole liver and 3 mm spinal cord tissues including the injured epicenter and 1.5 mm away from the injured epicenter on both the rostral and caudal sides were harvested from each mouse at various timepoints. The liver and spinal cord tissues were then processed into single-cell suspensions. Cells were stained with Zombie Violet™ FVD to exclude FVD-positive dead cells, followed by incubation with CD16/CD32 mAbs for blocking FcR binding, and fluorochrome-conjugated mAbs against mouse CD45, CD3, CD4, CD8, CD11b, Ly-6G, Ly-6C, F4/80, B220, and NK1.1. Appropriate isotype controls were used at the same protein concentrations as the test mAbs. Tissue-infiltrating leukocytes (LYM) were defined as CD45High cells. Within the CD45High population, polymorphonuclear neutrophils (PMNs) were identified by Ly-6G expression, T cells as CD45HighCD3+ cells, CD4 T cells as CD45+CD3+CD4+, CD8 T cells as CD45+CD3+CD8+, B cells as CD45+B220+, NKT cells (CD45+CD3+NK1.1+), and monocytes/macrophages (M/M*ϕ*) as CD45+CD11b+Ly-6G−. M/M*ϕ* were further divided into inflammatory (Ly-6CHigh ) and tissue-repairing (Ly-6CLowF4/80High) subsets. **A)** Gating strategy for flow cytometric analysis of immune cells in the spinal cord (upper panel) and livers (lower panel) of WT vs CD1dKO mice post-SCI. Gated M/Mϕ (M) showing the % of Ly-6CHigh versus Ly-6CLowF4/80High M/Mϕ in the spinal cord and liver. NKT cells (CD45+CD3+NK1.1+) were barely detected in the spinal cord from both WT and CD1dKO mice with and without a SCI (**A**, right panel). **B)** Pooled data of CD45+ lymphocytes, total T cells, CD4+ T cells, CD8+ T cells, B220+ B cells, and PMN in the spinal cord on days 1 (n=4/group), 3 (n=4/group), 7 (n=4/group) and 28 (n=4/group) post-SCI.

### Supplementary Figure 2

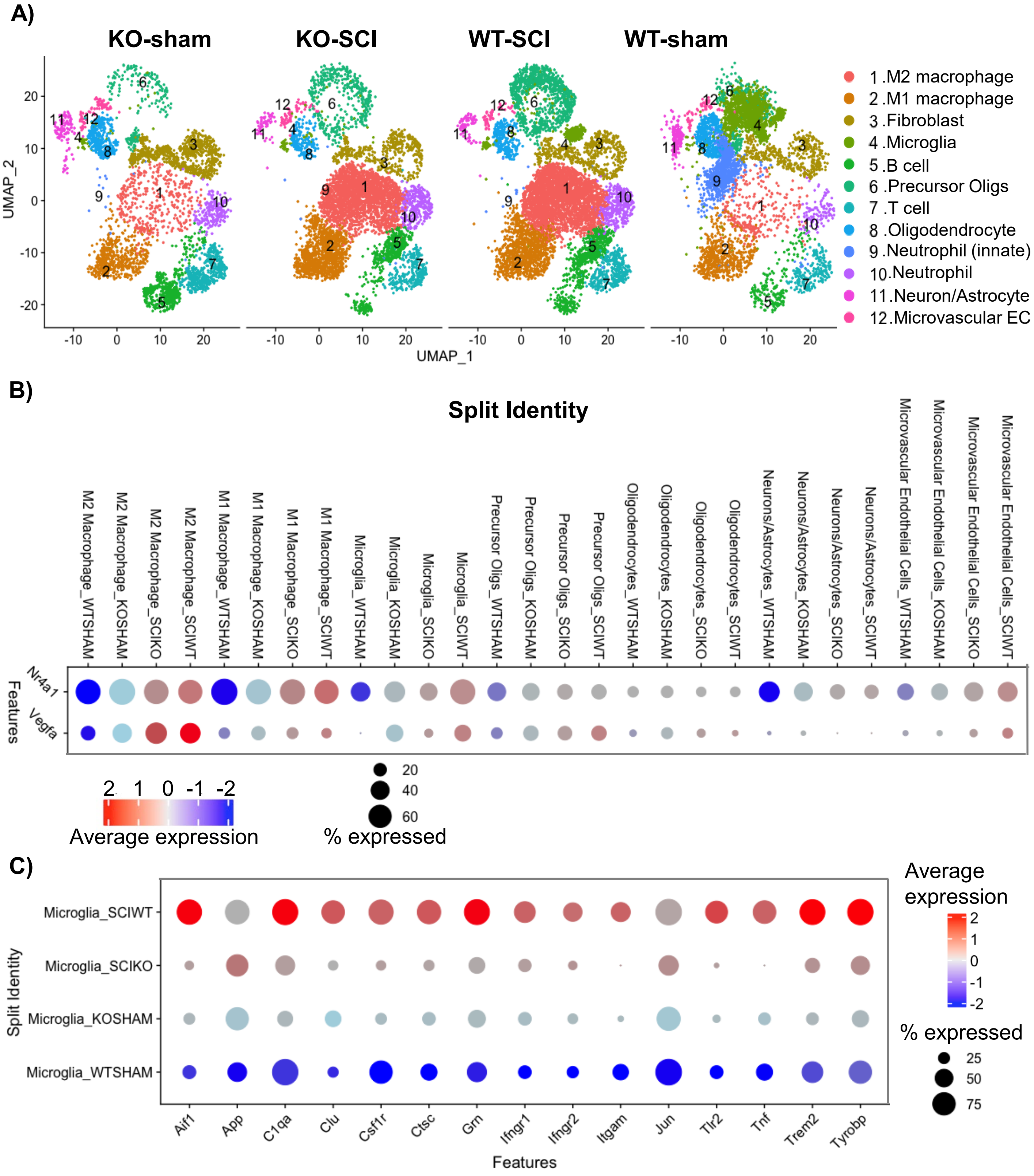

**Figure S2. scRNA-seq transcriptomics analysis of injured spinal cord cells.** WT and CD1dKO mice received either a SCI or sham injury. On day 7, a 2-cm section of spinal cord containing the injured site was used for isolating single cell populations for scRNA-seq analysis. **A)** UMAP plots showing 12 major cell types pooled from each group of mice. Each dot represents a single cell. **B)** Gene expression of *Vegfa* and *Nr4a1* in different cell types in spinal cord tissues including microvascular endothelial cells (ECs), neurons, astrocytes, oligodendrocytes (ODCs), precursor oligs, microglia, and macrophages (M1 and M2). **C**) Transcriptional signature of microglia activation. Clustering analysis was performed using R studio (version 3.6.1). Four mice were used per condition.

**Supplementary Table 1. Cytokine profiles in the spinal cord tissues from WT and CD1dKO mice after SCI (24 h post-SCI)**

| Cytokine | WT (pg/mL)) | KO (pg/mL) | *p* value | Cytokine | WT (pg/mL) | KO (pg/mL) | *p* value |
|---|---|---|---|---|---|---|---|
| **G-CSF** | 12846 ± 4420 | 7386 ± 5418 | 0.12 | **GM-CSF** | 42.2 ± 6.3 | 18.4 ± 7.3 | 0.15 |
| **IL-1β** | 19.1 ± 7.6 | 24.7 ± 6.1 | 0.17 | **IL-3** | UD | UD | |
| **IL-2** | 29.0 ± 5.6 | 23.9 ± 2.2 | 0.05 | **IL-5** | 7.7 ± 2.7 | 6.8 ± 2.4 | 0.52 |
| **IL-9** | 206.7 ± 56.5 | 177.3 ± 34.8 | 0.27 | **IL-12p70** | 8.9 ± 2.9 | 5.5 ± 2.9 | 0.053 |
| **IL-12p40** | 50.2 ± 9.7 | 41.0 ± 6.3 | 0.06 | **IL-13** | 94.7 ± 20.4 | 76.8 ± 21.3 | 0.15 |
| **LIF** | 201.0 ± 154.8 | 170.4 ± 100.9 | 0.68 | **IL-15** | 215.9 ± 39.2 | 180.3 ± 30.7 | 0.09 |
| **LIX** | 72.2 ± 28.1 | 46.3 ± 17.7 | 0.07 | **IL-10** | 18.7 ± 3.9 | 16.4 ± 4.6 | 0.21 |
| **IL-17** | 2.9 ± 1.0 | 2.4 ± 0.3 | 0.23 | **IP-10** | 862.3 ± 355.9 | 1030 ± 214 | 0.32 |
| **KC** | 5811 ± 2771 | 2764 ± 2593 | 0.07 | **MCP-1** | 2012 ± 731 | 1256 ± 824 | 0.11 |
| **MIP-1α** | 108.0 ± 11.1 | 113.8 ± 19.0 | 0.52 | **MIP-1β** | 36.4 ± 4.9 | 74.3 ± 38.8 | 0.044 |
| **M-CSF** | 67.2 ± 26.6 | 48.9 ± 12.5 | 0.13 | **MIP-2** | 4226 ± 2197 | 1932 ± 1309 | 0.07 |
| **MIG** | 17.0 ± 6.8 | 14.7 ± 5.4 | 0.51 | **RANTES** | 20.8 ± 4.2 | 23.4 ± 7.8 | 0.48 |

**Note:** WT (wild-type): n=6; OK (CD1dKO): n=7. All values: mean ± SD; UD, undetectable. Statistical analysis was performed using Student's *t*-test.
